# Capacity and tradeoffs in neural encoding of concurrent speech during Selective and Distributed Attention

**DOI:** 10.1101/2022.02.08.479628

**Authors:** Maya Kaufman, Elana Zion Golumbic

**Affiliations:** The Gonda Center for Multidisciplinary Brain Research, Bar Ilan University, Ramat Gan, Israel

## Abstract

Speech comprehension is severely compromised when several people talk at once, due to limited perceptual and cognitive resources. Under some circumstances listeners can employ top-down attention to prioritize the processing of task-relevant speech. However, whether the system can effectively represent more than one speech input remains highly debated.

Here we studied how task-relevance affects the neural representation of concurrent speakers under two extreme conditions: when only <u>one</u> speaker was task-relevant (Selective Attention), vs. when <u>two</u> speakers were equally relevant (Distributed Attention). Neural activity was measured using magnetoencephalography (MEG) and we analysed the speech-tracking responses to both speakers. Crucially, we explored different hypotheses as to how the brain may have represented the two speech streams, without making a-priori assumptions regarding participants’ internal allocation of attention.

Results indicate that neural tracking of concurrent speech did not fully mirror their instructed task-relevance. When Distributed Attention was required, we observed a tradeoff between the two speakers despite their equal task-relevance, akin to the top-down modulation observed during Selective Attention. This points to the system’s inherent limitation to fully process two speech streams, and highlights the complex nature of attention, particularly for continuous speech.

## Introduction

One of the hallmark signatures of attention on neural processing is the top-down modulation of activity in sensory cortices, whereby responses to stimuli that are in the focus of attention are amplified relative to behaviorally irrelevant stimuli (Hillyard et al. 1973; Hansen and Hillyard 1983; Woods et al. 1984; Bidet-Caulet et al. 2007; Manting et al. 2020). These modulatory effects are recognized as critical for gating the transmission of task-relevant input from sensory cortex to higher-order brain regions (Broadbent, 1958; Lachter, 2004). This early-gating is assumed to be particularly important in the case of speech-processing, since the language-system suffers from limited capacity for applying linguistic processing to multiple concurrent speech-stimuli (‘processing bottlenecks’; Treisman et al. 1964; wood and cowan 1995; Koelewijn et al. 2012; Bronkhorst, 2015; Mccloy and Lee 2015). In line with this assumption, a plethora of research in recent years has shown that the neural speech tracking response is significantly modulated by attention, with enhanced representation of to-be-attended speech observed in both auditory cortex and in language-processing regions (Kerlin et al. 2010; Ding and Simon 2012a; Ding and Simon 2012b; Mesgarani and Chang 2012; Power et al. 2012; Zion Golumbic et al. 2013; O’Sullivan et al. 2015; Fuglsang et al. 2017; Fiedler et al. 2019).

And yet, these modulatory top-down effects do not eliminate the internal representation of task-irrelevant (or ‘to-be-ignored’) speech. Not only is this speech still encoded in auditory cortex (Ding and Simon 2012a; Horton and Srinivasan 2013; Zion Golumbic et al. 2013; O’Sullivan et al. 2015; Fiedler et al. 2019), but a multitude of behavioural and neural findings provide evidence that some linguistic processing is also applied to task-irrelevant speech (Tun et al. 2002; Dupoux et al. 2003; Beaman et al. 2007; Rivenez et al. 2008; Carey et al. 2014; Parmentier et al. 2014, 2018; Schepman et al. 2015; Vachon et al. 2019; Dai et al. 2021; Har-shai Yahav and Zion Golumbic, 2021). Therefore, alongside the convincing evidence that selective-attention *biases* neural processing such that to-be-attended speech is preferentially encoded, we still do not have a full understanding of the system’s capacity and limitations for processing additional concurrent speech.

One reason it has been difficult to reach a consensus about whether linguistic processing is applied to task-irrelevant speech in Selective Attention ‘cocktail party’ paradigms is the inherent ambiguity regarding participants’ internal focus of attention: We *cannot know for sure* whether individuals actually focused their attention exclusively towards the designated speaker. In reality, it is highly possible that, even when instructed to pay attention to only one speaker, individuals occasionally switch their attention to the other speaker, or just listen to both of them, especially when presented with long narratives of continuous speech. Indeed, many of the insertion and priming effects observed for task-irrelevant speech have been explained by some as resulting from momentary attention switches between streams, rather than an indication of true parallel processing (wood and cowan 1995; Dupoux et al. 2003; Escera et al. 2003; Lachter 2004; Beaman et al. 2007). This ‘ground-truth’ problem is extremely difficult to address empirically, since we lack a reliable readout of an individual’s internal allocation of attention. Unfortunately, most behavioural and neural measures used in in Selective Attention paradigms cannot provide sufficient temporal resolution to indicate the momentary locus of attention and/or how well task-irrelevant speech has been processed (Har-shai Yahav and Zion Golumbic 2021). While some attempts have been made at utilizing advanced analytical approaches to achieve real-time measures of attention (Miran et al. 2018; Jaeger et al. 2020), these studies are still limited by the lack of access to the true internal state of attention.

Given this inherent limitation, in the current study we take a different approach to studying the co-representation of competing speech in the brain and ask: to what extent is the system *capable* of processing, and ultimately comprehending, two narratives presented concurrently? Specifically, in this magnetoencephalography (MEG) study we compared speech comprehension and neural tracking of concurrent speech under two extreme conditions: the conventional Selective Attention task, where only one speaker is designated ‘to-be-attended’, and a Distributed Attention task, in which participants are asked to listen to and comprehend both speakers. The Distributed Attention condition encourages participants to optimize their listening strategy and to actively try to pick up information from both streams at once. Hence, it offers a window into the system’s capacity and limitation for processing two concurrent speech streams and provides the opportunity to estimate potential tradeoffs in their processing. Specifically, it allowed us to contrast two opposing hypotheses regarding the co-representation of competing speech: Are both speakers comprehended and represented in the brain (‘equal-representation’ hypothesis; Deutsch and Deutsch 1963; Parmentier 2008; Parmentier et al. 2018; Vachon et al. 2019)? Or, is the system only capable of processing one speech-stream at a time, resulting in a tradeoff in the neural representation of concurrent speech during Distributed Attention (‘biased-representation’ hypothesis; Buschman and Miller 2010; Vestergaard et al. 2011; Larson and Lee 2013; Hambrook and Tata 2019; Huang and Elhilali 2020)?

Importantly, when analysing the speech tracking response in the Selective and Distributed attention conditions, we refrained from making a-priori assumptions about how individuals actually allocated their attention. Rather, we used a data-driven approach to examine different potential listening strategies and test which ones best explain the recorded neural data, at the level of individual participants. By investigating both Selective and Distributed Attention, this study provides insight into the interaction between instructed task-relevance and internal allocation of resources for representing and processing concurrent speech.

## Methods

### Participants

33 Hebrew native speakers, right-handed participants (19 female, mean age 24.5 ± 5.9, range 18-48), with self-reported normal hearing and no history of neuropsychiatric disorders took part in this study (one participant reported daily anti-depressants use). Five participants were excluded from the data analysis due to excessive artifacts in the MEG data. One additional participant was excluded from the data analysis due to credibility issues. The study was approved by the Institutional Ethics Committee at Bar Ilan University. Participants provided their written consent prior to commencement of the experiment and were either paid or received academic credit points for their participation.

### Natural speech stimuli

Speech stimuli consisted of short personal narratives (42-46 sec long) pre-recorded by two actors (one female, one male). In order to preserve the natural speech characteristics of the speech, the recordings were only minimally edited using the softwares Praat (http://www.praat.org, version 6.0.37) and Audacity (https://www.audacityteam.org, version 2.1.3) to equate the perceived sound loudness of all recordings. Sound volume was adjusted separately for the female and male speakers, to account for the perceptual amplification of high-pitch speech.

### Experimental procedure

In each trial, participants were presented with a pair of speech-stimuli – one by the female and one by the male speaker – presented to different ears (dichotically). Before each trial, participants were either instructed to attend to one speaker (Selective Attention condition) or to both speakers (Distributed Attention condition), using a visual cue (Fig. 1). The visual cue remained on the screen throughout the trial, serving as a reminder for the instructions. In the Selective Attention condition, following each trial participants answered three multiple-choice comprehension questions about the designated to-be-attended narrative (chance level = 25%). In the Distributed Attention condition, three consecutive comprehension questions were asked about each of the narratives (the order of presentation for the two sets of questions was equally balanced between the female/male narrative and between the right/left ear). The questions could be either about specific details in the narrative (e.g., “what was the name of the speaker’s daughter?”), or could refer to a broad concept in the narrative (e.g. “what was the speaker excited about?”).

**Figure 1:**
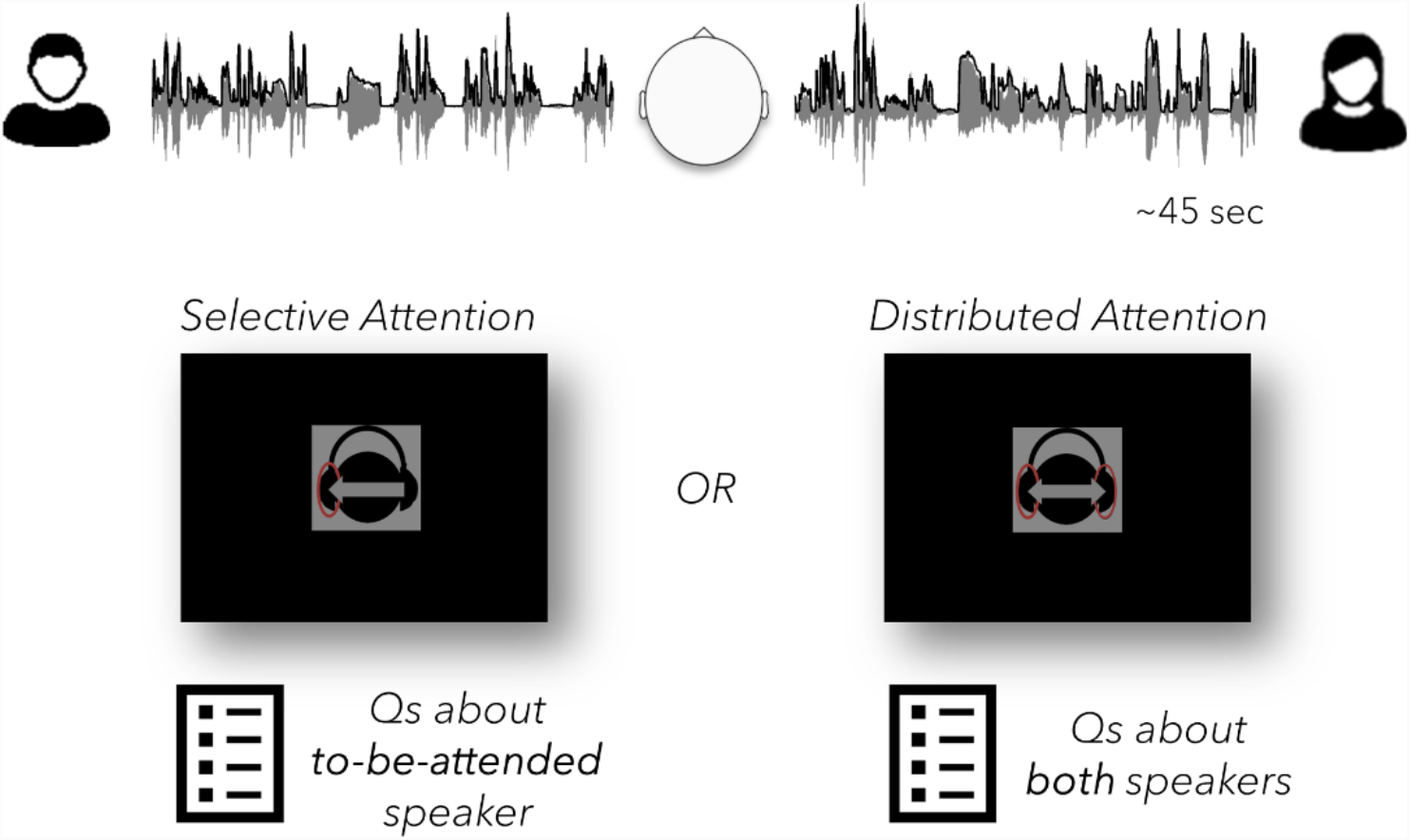
Illustration of the experimental paradigm. In each trial, a pair of natural Hebrew speech narratives (one female, one male voice) were presented dichotically through earphones. The spatial location of each voice (right/left) was counter-balanced across trials. Before each trial, participants were instructed by a visual cue to either attend to one of the speakers (Selective Attention condition) or to both speakers (Distributed Attention condition). Following each trial, multiple choice questions were asked about the content of the to-be-attended narrative (Selective Attention) or about both narratives (Distributed Attention).

**Figure 2:**
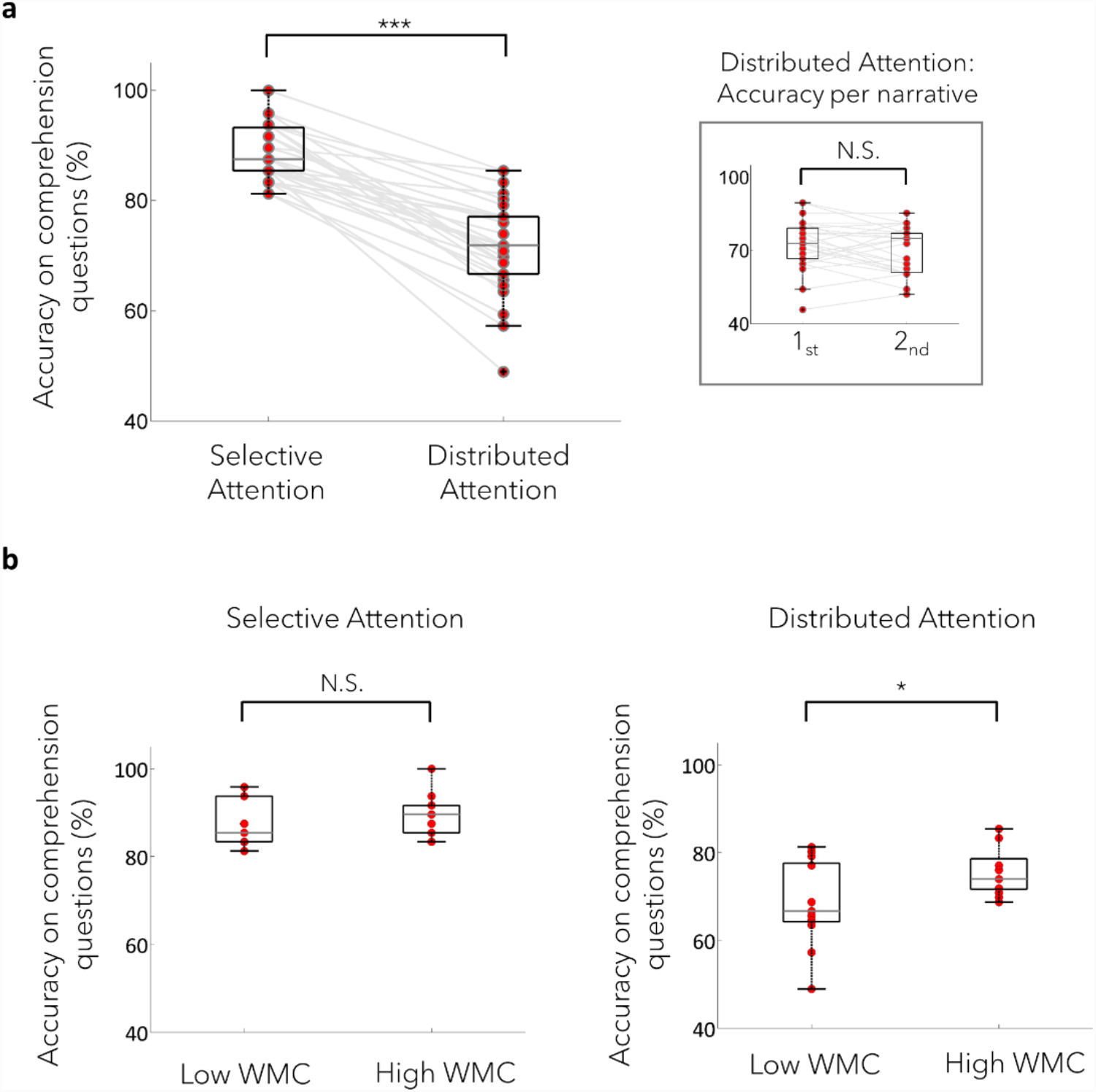
Performance on the behavioural task. **(a)** Mean accuracy on comprehension questions across all participants (n=27) in the Selective vs. Distributed Attention condition. Asterix represent statistical significance between the two conditions (p<10^−10^), with significantly better comprehension for the Selective Attention condition. Right inset: Mean accuracy on the comprehension questions in the Distributed Attention condition, separately for the 1^st^ and 2^nd^ set of questions asked. Lines represent individual results and boxes represent the 25-75^%tile^, in both the main figure and inset. **(b)** Correlation between working memory capacitance (WMC) and behavioural performance. Asterix represent statistical significance on the median split, indicating significantly better performance for participants with high vs. low WMC on the Distributed Attention task (p=0.02), but not on the Selective Attention task (p=0.5).

The pairing of female-male speech stimuli remained constant across all participants, however the assignment of each pair to an attentional condition (Selective/Distributed), their order of presentation in the experiment and the status of each narrative as to-be-attended or task-irrelevant in the Selective Attention condition was pseudo-randomized across participants. The lateralization of speaker presentation and the instructed direction of attention in the Selective attention condition (left/right ear) were also counter-balanced across conditions within subject.

Accordingly, the female speaker was presented to the left ear in exactly half of the trials, of which half belonged to the Distributed attention condition, and half to the Selective attention condition. Additionally, in the Selective Attention condition, the instructed direction of attention was left in half of the trials, of which half were attend-female and half attend-male trials (see illustrations in Figures 3a and 5a). The experiment consisted of a total of 32 trials, 16 per attention condition. Before starting the experiment, participants were familiarized with the stimuli and task in a training trial. The total duration of the experiment was 47.76±5.03 (range 41.19-65.42) minutes.

**Figure 3:**
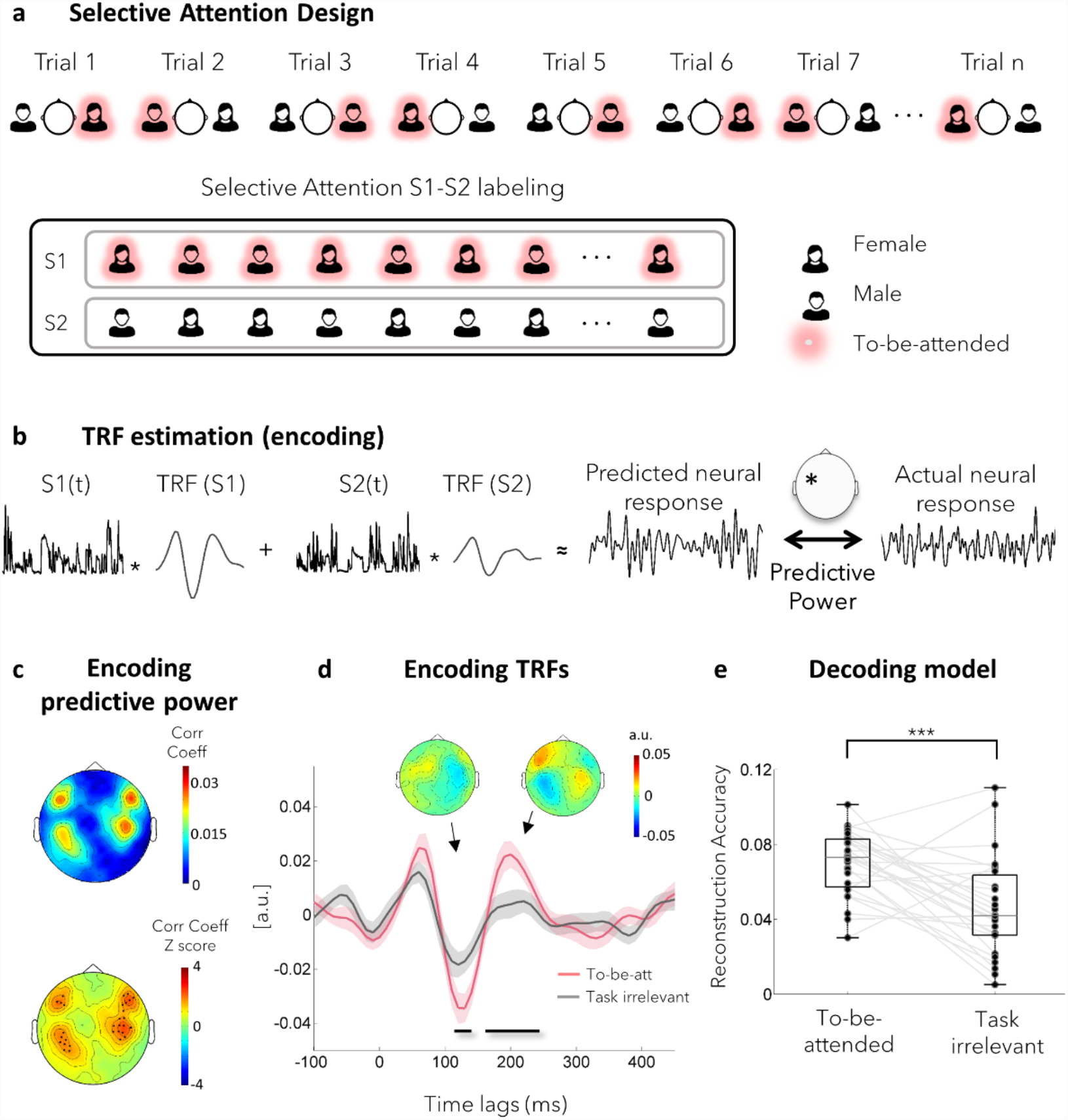
Selective Attention: Group level results. **(a)** Top panel: Selective Attention experimental design. In each trial, one of the speakers was pre-designated as the to-be-attended speaker (highlighted in pink). The identity (female/male) and presentation ear (left/right) of the to-be-attended speaker was counter-balanced across trials. Bottom panel: Illustration of the S1-S2 labeling scheme in the Selective Attention condition. The narratives designated as to-be-attended from all trials were labeled as S1, and the task-irrelevant narratives were labeled as S2. This labeling scheme is used to create two regressors, consisting of the speech-envelopes of the narratives included in S1 and S2 respectively. **(b)** Illustration of the encoding speech tracking rationale. Separate TRFs are optimized for each regressor (S1, S2) at each sensor, so as to maximize the correlation between the predicted neural response of the model and the actual recorded neural response (predictive power). **(c)** Topography of the predictive power (top) and normalized predictive power (relative to a null-distribution of permuted data, bottom) of the encoding model, averaged across participants. Both topographies present the common four-pole topography characteristic of auditory responses. The MEG sensors marked with an asterix in the bottom panel are those where the normalized predictive power significantly exceeded chance level (Z>1.64), which is indicative of significant speech tracking. These sensors were used as a region of interest (ROI) for further analysis. **(d)** The resulting TRFs for to-be-attended (pink) and task-irrelevant (grey) speech, averaged across participants, from the sensor with the highest normalized predictive power. Shaded highlights denote SEM across participants. Horizontal black lines represent the time-windows with statistically significant differences between the TRFs (120-140ms and 160-240ms; p<0.002, corrected). The topographies of the differences between the TRFs in each time window are presented above the waveforms. **(e)** Reconstruction accuracy of the two speech stimuli across all participants, using the decoding approach. Lines represent individual results and boxes represent the 25-75^%tile^.

### Working Memory test

Participants’ Working Memory capacity (WMC) was evaluated using the Operation Span test (OSPAN version 2.0.10, Foster et al. 2015, adaptation to Hebrew taken from https://englelab.gatech.edu/translatedtasks.html#hebrew). In this task participants are required to recall a set of 5 Hebrew letters while solving simple arithmetic problems. The test was comprised of 3 blocks, with 25 arithmetic problems per block (total duration was 14.27-3.18, range 9.48-19.68 minutes). The test produces two main scores referring to the number of letters recalled and positioned correctly within the sequence (1) including only trials where all letters were correctly recalled (absolute OSPAN score) and (2) including all trials (partial OSPAN score). However, we used only the partial scores, as per recommendation of previous studies, since they make use of all available information (Friedman and Miyake 2005; Redick et al. 2012; Ðokić et al. 2018).

### MEG data acquisition

MEG data was acquired in supine position, using a whole-head 248-channel magnetometer array (4-D Neuroimaging, Magnes 3600 WH) in a dimly and magnetically shielded room. Data were sampled at 1017.23 Hz and bandpass-filtered online at 0.1 to 400 Hz. Head position in relation to the MEG sensors was estimated using 5 coils that were attached to the participant’s scalp. Head shape was digitized manually for later source estimation. Speech stimuli were presented to both ears via MEG compatible in-ear tubephones (ER13-14, Etymotic Research), at a comfortable listening level that remained constant for all participants.

### Data analysis

#### Behavior analysis

Accuracy levels were defined as the percentage of correct answers out of the total number of comprehension questions asked (3 per trial in the Selective condition and 6 per trial in the Distributed condition). Differences in accuracy in the Selective and Distributed Attention conditions were evaluated using a paired t-test. In the Distributed Attention condition we additionally compared accuracy on the first vs. second set of questions asked (about each of the two narratives) using a paired t-test, in order to test for potential primacy effects, and to make the accuracy levels more comparable to the Selective Attention condition, where only three questions were asked. Since previous studies have suggested that working memory capacity (WMC) is positively correlated with enhanced attention capacity, we additionally tested whether behavior on either task was better in participants with high vs. low WMC using a Wilcoxon ranksum test (based on a median-split of WMC scores; OSPAN test, partial score).

#### MEG data pre-processing

MEG data pre-processing was performed in MATLAB (R2013b, The MathWorks) using the FieldTrip toolbox (Oostenveld et al. 2011) and custom written scripts. External noise (power-line, mechanical vibrations of the building), as well as heart-beat artifacts, were removed offline using a predesigned algorithm (Tal and Abeles 2013). Independent component analysis (ICA) was used for removal of blinks, eye movements and residual heartbeat traces. Two consistently noisy sensors were completely removed from the MEG neural data, such that a total of 246 sensors remained. No additional artifact rejection was performed. The cleaned neural data were segmented into trials, starting from ∼10 sec after the onset of the auditory stimuli (to avoid introducing onset effects and to allow sufficient time to adapt to the type of attention required), and until the offset of the shorter narrative in the presented pair. The segmented data were bandpass filtered between 1-20 Hz using a zero-phase FIR filter, and then downsampled to 100 Hz, for computational efficiency.

#### Speech Tracking Analysis: General

To estimate the neural response to the two speakers we performed speech tracking analysis, using both an encoding and a decoding approach. Briefly, in this analysis a Temporal Response Function (TRF) is estimated, which constitutes a linear transfer function describing the relationship between feature(s) of the stimulus (S) and the neural response (R) recorded when hearing it. Since in the current study two narratives were presented concurrently, we applied a multivariate approach to estimate the neural responses to both speakers (denoted as S1 and S2) using the mTRF MATLAB toolbox (Crosse et al. 2016). Specifically we used the broadband envelopes of the speech as stimulus features, which were extracted using an equally spaced filterbank between 100 to 10000 Hz based on Liberman’s cochlear frequency map. The narrowband filtered signals were summed across bands after taking the absolute value of the Hilbert transform for each one, resulting in a broadband envelope signal. This envelope was further downsampled to 100Hz and the first 10 seconds of each stimulus was removed, to match the MEG data.

In the encoding approach for speech tracking analysis, two separate TRFs are estimated per MEG sensor. These reflect transfer-function responses to each of the two speakers (S1 and S2), optimized so that their linear combination best predicts the neural response (Figure 3b). These TRFs can be thought of as a linear composition of partially overlapping neural responses to a continuous stimulus and are therefore conceptually analogous to Event Related Potentials (ERPs). As such, the peaks and throughs of the TRF can be interpreted in a similar fashion as more traditional ERP components. Hence, the advantage of encoding models is that they provide a detailed description of the neural response to each stimulus, by describing the time-course and topography of the response across sensors.

A complementary way to examine the neural representation of continuous speech is the decoding approach. In the decoding model a multidimensional transfer function is estimated using the neural data from all MEG sensors as input (R) in attempt to reconstruct the stimuli (S1 and S2). The advantage of the decoding model is the simplicity of its results, which gives a single value for each stimulus – the Pearson correlation between the reconstructed stimulus and the real stimulus. Thus the decoding benefits from simpler statistical analysis, whereas the results of the encoding give a more detailed description of the neural response but also require correction for multiple comparisons (across time and sensors). In the current project we used both encoding and decoding approaches, utilizing the advantages of each one.

Encoding and decoding models were optimized separately for the Selective and Distributed conditions. Encoding TRFs were calculated over time lags ranging from -150 (pre-stimulus) to 450 msec, and the decoding analysis used time lags of -400 to 0 msec. Both models were optimized using a leave-one-out approach, where in each iteration, 15 trials are selected to train the model (train set), which was then used to predict either the neural response at each sensor (encoding) or the speech envelope of the two speakers (decoding) in the left-out trial (test set). The model’s goodness of fit (predictive power) is the Pearson correlation between the predicted and the actual signal. This procedure is repeated 16 times, with a different train-test partition in each iteration, and the predictive power is averaged over all 16 iterations. To prevent overfitting of the model, a ridge parameter (λ) was chosen as part of the cross-validation process. This parameter significantly influences the shape and amplitude of the TRF and therefore, rather than choosing a different λ for each participant (which would limit group-level analyses), a common λ value was selected for all participants. Specifically, for each participant, λ values ranging from 2^-5^ to 2^15^ were tested and the value yielding the average highest predictive power (across all channels\participants) was chosen as the common optimal λ. For the encoding model this value was λ=2^5^ and for the decoding model λ = 2^9^. All results reported in here use these ridge parameters.

Before testing for effects of attention, we first determined which subset of sensors showed a significant speech tracking response. This was done using a permutation test, where we shuffled the pairing between acoustic envelopes (S1 and S2) and neural data responses (R) across trials such that speech-envelopes presented in one trial were paired with the neural response recorded in a different trial. This procedure was repeated 100 times and an encoding model was optimized for each permutation, yielding a null-distribution of predictive power values that could be obtained by chance for each MEG sensor. We then z-scored the predictive power values from the true pairing relative to this null-distribution. The sensors that exceeded an average of Z > 1.64 across participants (one-sided, p<0.05) were considered as displaying a significant speech tracking neural response and were used as a Region of Interest (ROI) for subsequent encoding analyses (in both the Selective and Distributed conditions).

#### Speech Tracking Analysis: Selective Attention condition

We next turned to evaluate the effects of attention on the speech tracking response. In the Selective Attention condition the instructions were very clear – attend to the designated speaker in each trial in order to answer questions about that narrative. Therefore, labeling the two speech-stimuli in each trial as S1 and S2 for the purpose of multivariate TRF analysis is straightforward: The to-be-attended speech in each trial was labeled as S1, and the task-irrelevant speech was labeled as S2. Note that since the gender and spatial location of the to-be-attended speech were fully counterbalanced across trials, the labels ‘S1’ and ‘S2’ do not represent specific speaker identities/ears, but only their designated attentional status. These regressors were used for both the encoding and decoding analyses.

The encoding model yields separate TRFs for to-be-attended and task-irrelevant speech at each MEG sensor (Figure 3b, in past studies these have been shown to be modulated by their designated attentional-status: Ding and Simon 2012a, 2012b; Power et al. 2012; Zion Golumbic et al. 2013; Fuglsang et al. 2017; Fiedler et al. 2019). To detect time-windows where these differences were statistically significant, we used a paired-comparison test, focusing on TRFs in the significant ROI, and using a permutations test to correct for multiple comparison across time-points. In the decoding approach, reconstruction accuracies are obtained for the to-be-attended and task-irrelevant, reflecting how well each of the two speech-envelopes could be estimated from the neural signal. These reconstruction accuracies were compared using a paired t-test.

In addition to these group-level analyses, we also wanted to assess to what degree the well-documented effect of Selective Attention was present in individual participants. This analysis used only the decoding approach, since it simplifies the comparison between S1 and S2 to a comparison of two reconstruction values, rather than including multiple comparisons over sensors and time. For each participant we calculated an Attention-Bias Index, which is the difference in the reconstruction accuracy for the to-be-attended and the task-irrelevant speakers. To account for individual differences in SNR and to obtain a data-drive estimation of chance-level, we used permutations to create an “**attention-agnostic”** null distribution of bias indices (illustrated in Fig. 4a). Specifically, in each permutation the two narratives were randomly re-labeled as S1/S2 so that the speech included in each regressor was 50% to-be-attended and 50% task-irrelevant. Two decoders were trained on the data using these attention-agnostic labelings and the reconstruction accuracy for each regressor was estimated as well as the difference between them (attention-agnostic bias index). We repeated this process 100 times, which resulted in a distribution of attention-agnostic bias indices that reflect the range of bias index values that could be obtained by-chance. The real Attention-Bias Index of each participant was normalized (z-scored) relative to this null-distribution. If the Selective Attention instructions truly lead to enhanced neural representation of the to-be-attended speaker relative to the task-irrelevant one, this would be reflected in a positive normalized Attention-Bias Index that exceeded Z = 1.96 (p<0.05, two-sided). However, if the normalized Attention-Bias Index falls within this null-distribution (Z < 1.96), would suggest that the difference between to-be-attended and task-irrelevant speech is not larger than could be achieved by chance.

**Figure 4.**
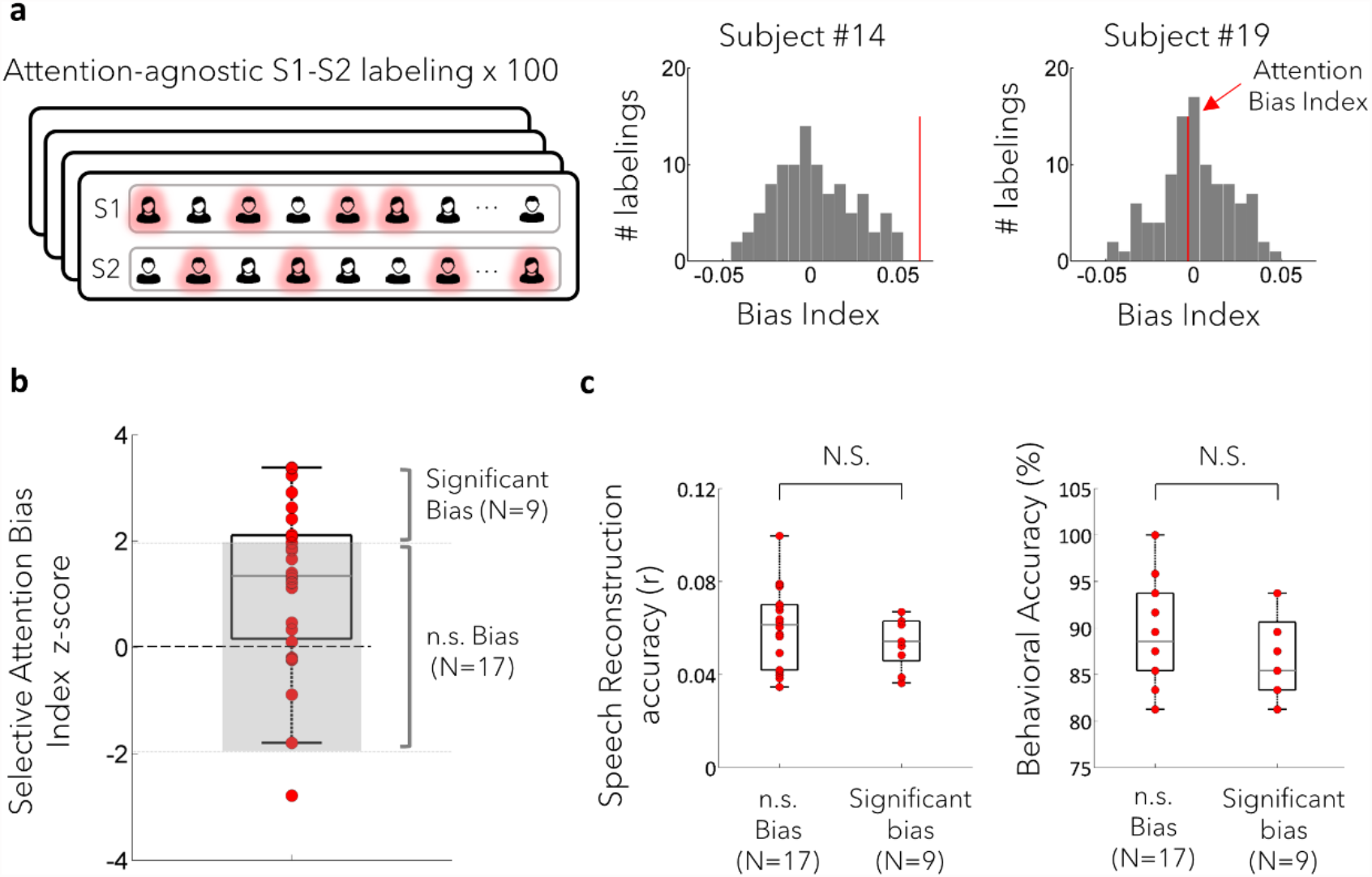
Selective Attention: Attention-bias in individual participants. (**a**) Illustration of the Attention-agnostic labeling scheme used to normalize the Attention-Bias Index. 100 S1-S2 labeling schemes were used, in which the to-be-attended narratives (highlighted in pink) were equally (and randomly) distributed between S1 and S2. A decoding analysis was performed for each labeling in which reconstruction accuracies were estimated for S1 and S2, and the difference between them calculated (Bias Index). This resulted in an attention-agnostic distribution of bias indices for each participant (two examples shown on the right). The true Attention-Bias Index, which was derived using the Selective Attention S1-S2 labeling (shown in Figure 3a, and as a red line in the two examples here), was compared and normalized relative to this null distribution. As shown in the two examples here, for some subjects the Attention-Bias Index was significantly higher than most/all attention-agnostic labelings (e.g. subject #14), indicating a significant bias of the neural speech tracking response towards the to-be-attended narrative. However, in others it fell well within the attention-agnostic null-distribution (e.g. subject #19), indicating no significant modulation by attention. (**b**) The normalized Selective Attention-Bias Index for all participants. The grey-shaded area represents chance-level bias values (−1.96 < Z < 1.96), and the majority of participants fall within this range, with only 30% (n=9) of participants displaying normalized Attention-Bias Index values that exceed chance. Red circles denote individual values and boxes represent the 25-75^%tile^. (**c**) Comparison of behaviour performance (right) and of the overall speech-reconstruction accuracies (averaged for both to-be-attended and task-irrelevant speakers; left) between the subgroup of participants who showed significant Attention-Bias (n=9) vs. those who did not (n=17). No significant differences between the subgroups were found for either measure. Red circles denote individual values and boxes represent the 25-75^%tile.^

#### Speech Tracking Analysis: Distributed Attention condition – testing systematic bias

Unlike the Selective Attention condition, in the Distributed Attention condition there is no a-priori “correct” way to label the two concurrent speakers as S1 and S2, since both speakers are equally task relevant. Instead, here we analyzed the data using several different labeling-schemes, which allowed us to test specific a-priori

hypotheses about how the brain represents two equally-task-relevant speakers. Specially, we wanted to test whether there is evidence that individuals still show a bias towards one of the speakers, despite their equal task-relevance. In our first analysis we tested for evidence for systematic-bias, manifesting either as a consistent preference of a particular speaker (gender-bias) or towards a particular ear (spatial-bias).

To test for ***systematic gender-bias***, we trained two decoders on a S1-S2 labeling scheme in which the audio from the female and male speakers in each trial were labeled as S1 and S2, respectively (Figure 5b, top panel). The reconstruction accuracies of the female and male speakers were estimated, and the differences between them was taken as the Gender-Bias Index, for each participant. A similar approach was used to test for ***systematic spatial-bias***, where two decoders were trained using a S1-S2 labeling scheme in which the audio presented from the left and right ears in each trial were respectively labeled as S1 and S2 (Figure 5c, top panel). Here too, we extracted reconstruction accuracies for each ear separately and calculated a Spatial-Bias Index, for each participant. Statistical evaluation of the Gender-Bias and Spatial-Bias at the group level was performed using a t-test vs. 0.

**Figure 5.**
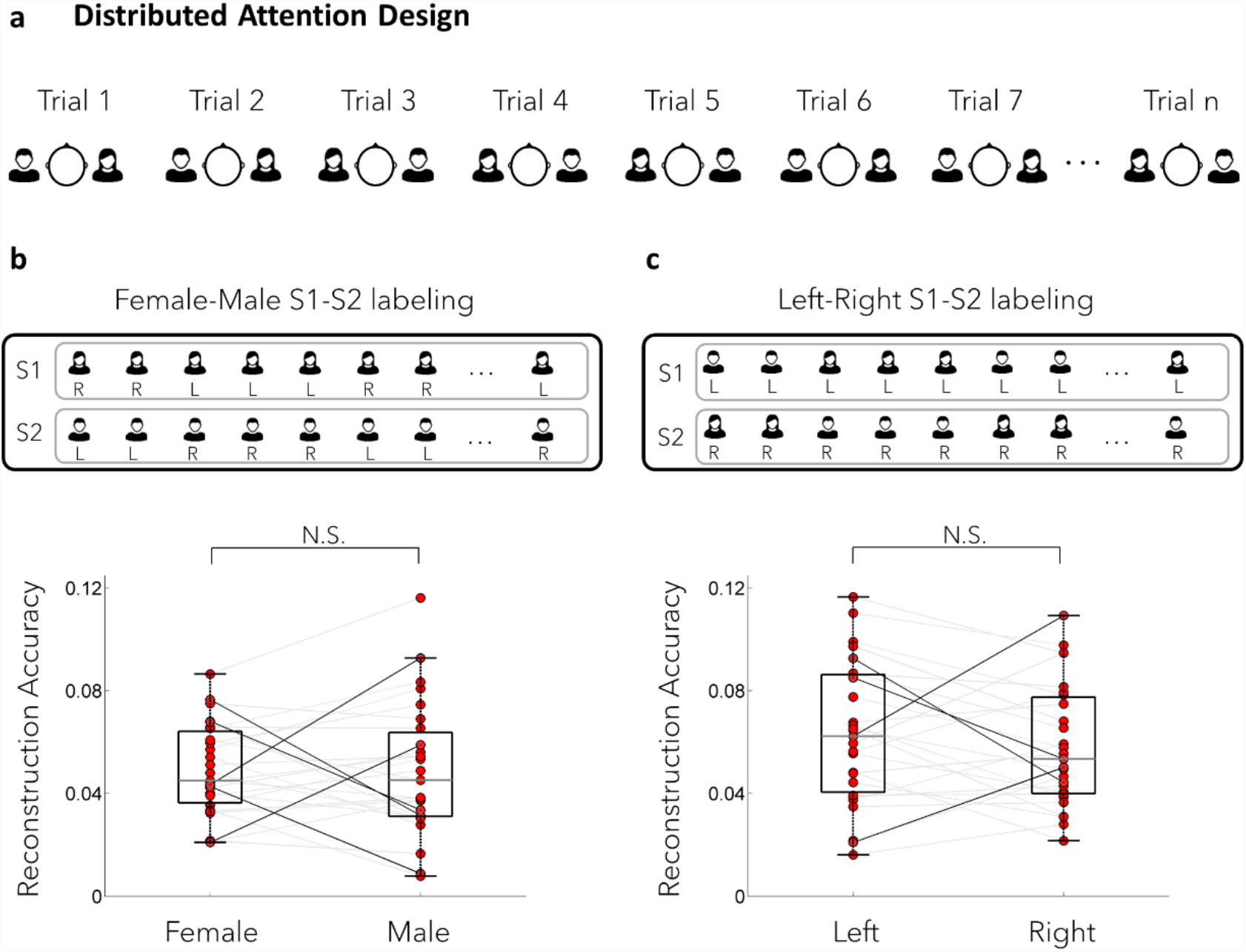
Distributed Attention: Systematic Bias. **(a)** Distributed Attention experimental design. In the Distributed Attention condition, both speakers were equally task-relevant. The allocation of each speaker to the left/right ear was counter-balanced across trials. **(b)** Investigation of systematic Gender-Bias. Top panel: To test for systematic Gender-bias, a S1-S2 labeling scheme was used in which the narratives of the female and male speakers were labeled as S1 and S2, respectively. Bottom: The reconstruction accuracies for the female and male regressors, for all participants. Lines represent individual values and boxes represent the 25-75^%tile^. Bold lines represent participants with statistically significant systematic bias towards one speaker (Z>1.96 or Z<-1.96, normalized against a gender-agnostic distribution). At the group level, there was no significant difference between the reconstruction accuracy of the female vs. male narratives (p=0.94). **(c)** Investigation of systematic Spatial-Bias. Top panel: To test for systematic Spatial-bias, a S1-S2 labeling scheme was used in which the narratives presented to the left and right ear were labeled as S1 and S2, respectively. Bottom: The reconstruction accuracies for the left and right regressors, for all participants. Lines represent individual values and boxes represent the 25-75^%tile^. Bold lines represent participants with statistically significant systematic bias towards one ear (Z>1.96 or Z<-1.96, normalized against a spatial-agnostic distribution). At the group level, there was no significant difference between the reconstruction accuracy of the narratives presented to the left vs. right ear (p=0.31).

Since systematic bias patterns likely differ across individuals, in addition to the group-level analysis, we also tested these biases at the individual level. To this end, the Gender-and Spatial-Bias indices of each participant were normalized (z-scored) relative to a distribution of bias-values obtained from 100 random S1-S2 labeling, where both the gender and spatial location were equally represented in both S1 and S2. We extracted the difference between S1 and S2 decoding accuracies from each random labeling, yielding a distribution of gender-agnostic and spatial-agnostic bias-indices. The true gender- and spatial-bias indices from each participant were normalized to this distribution, and were considered significant if normalized values exceeded Z = ±1.96 (p<0.05, two sided). *Speech Tracking Analysis: Distributed Attention condition – searchlight approach for testing non-systematic bias*

Besides the possibility of systematic-bias towards one speaker/ear, another hypothesis is that listeners favor a different narrative in each trial in a ***non-systemati*c** fashion. Such non-systematic bias could be driven, for example, by the salience of the narrative-content, personal interest, or other features of the narrative. Uncovering this type of bias is extremely complicated, since we do not have empirical access to the ground-truth of participants’ internal preferences. To overcome this uncertainty, here we employ a searchlight approach, which allowed us to evaluate whether the recorded neural activity in the Distributed Attention condition fits with the predictions of the **equal-representation hypothesis**, according to which both speakers are equally represented in the brain, or whether there is evidence for a tradeoff between the neural representation of two speakers, in line with the **biased-representation hypothesis**.

The rationale guiding the searchlight approach is illustrated in Figure 6, and is as follows: For each participant we created 100 different versions of S1-S2 labelings, randomly assigning the two narratives in each trial either to S1 or to S2 (Fig. 6a). Under the **biased-representation hypothesis**, which posits that one speaker is prioritized in each trial, each S1-S2 labeling scheme represents a hypothetical listening/prioritization strategy that the listener could have adopted, in which one speaker in each trial is preferred relative to the other. Even though we do not have direct access into the ‘true’ internal prioritization, by testing multiple S1-S2 labelings some of them are likely to have a higher prevalence of the ‘true’ prioritized-speakers from different trials in one of the regressors. This maps onto specific predictions for how the predictive power (encoding) and reconstruction accuracy (decoding) would vary across S1-S2 labelings. Specifically, in the encoding analysis we would expect that S1-S2 labelings which have the highest predictive power (i.e., explain more of the variance in the neural response), also result in larger differences in the TRF peaks for S1 and S2, akin to what is traditionally found for Selective Attention (Fig. 6b, left panel). Similarly, in the decoding analysis, we would expect to find a tradeoff in reconstruction accuracy between the S1 and S2, with the labelings that yield high

**Figure 6.**
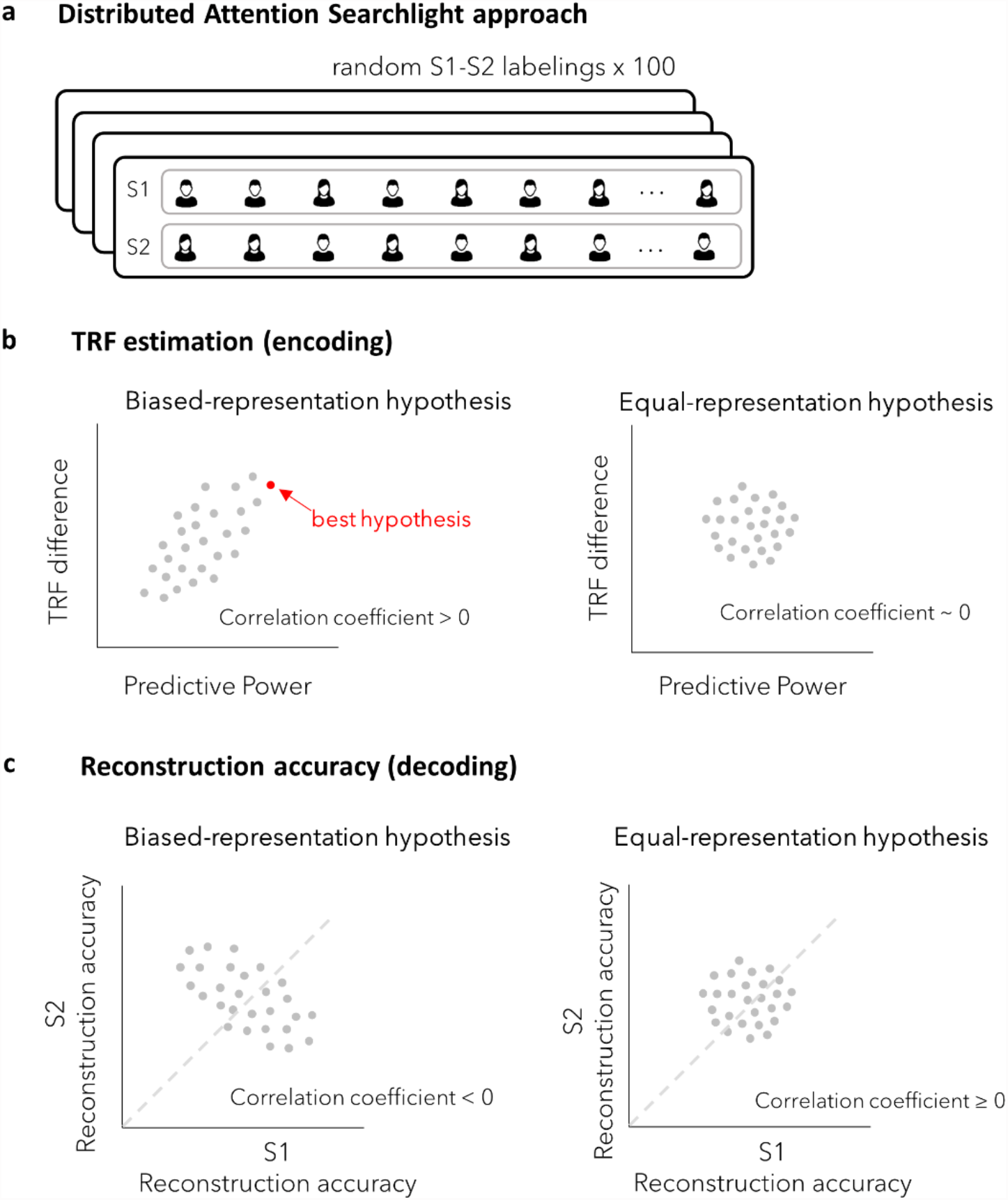
Distributed Attention: Searchlight approach rationale. **(a)** Illustration of the searchlight approach employed to study speech tracking under Distributed Attention. In each of 100 labelings, the two narratives in each trial were randomly assigned to S1 and S2. Both encoding and decoding speech tracking analyses were applied to each of these labeling-schemes, and the resulting distributions were used for testing the predictions of the equal vs. biased representation hypotheses. **(b)** Theoretical predictions for the equal-representation (right) and biased-representation (left) hypotheses, for encoding results: Under the **biased-representation hypothesis**, we would expect a positive correlation between the difference in the TRFs estimated for S1 and S2 and the predictive power of the model, with the labeling schemes that best estimates the neural response (i.e., with the highest predictive power; ‘best hypothesis’ – indicated by a red dot) to also have the highest difference between the two estimated TRFs. Conversely under the **equal-representation hypothesis**, we would expect both narratives to be similarly represented in the neural response and therefore any differences between the TRFs are not expected to produce better predictions of the neural response. This would manifest in no significant correlation. **(c)** Theoretical predictions for the equal-representation (right) and biased-representation (left) hypotheses, for decoding results: Under the **biased-representation hypothesis**, we expect the reconstruction accuracies of the two streams to demonstrate a tradeoff, manifested in a negative correlation between the reconstruction of S1 and S2. Conversely, under the **equal-representation hypothesis**, we do not expect to find any correlation between the reconstruction accuracies of the two streams (or perhaps a positive correlation), as they are expected to be similarly represented in the neural response.

reconstruction accuracy for one regressor associated with low reconstruction accuracy for the other (Fig. 6C, left panel). This pattern would highly support the biased-representation hypothesis. Conversely, under the **equal-representation hypothesis**, the allocation of speakers to S1/S2 should be fully interchangeable and therefore no significant differences would be expected for different S1-S2 labelings. Specifically, in the encoding analysis, the predictive power obtained for different S1-S2 labelings should not be systematically associated with differences in the estimated TRFs for the two regressors (Fig. 6b, right panel). Moreover, in the decoding analysis, reconstruction accuracy should be similar for S1 and S2, regardless of which narrative was assigned to each regressor, and there should be either a positive or null correlation between them (Fig. 6c, right panel).

To test these predictions, we conducted encoding and decoding analysis of the data, using 100 different S1-S2 lablelings. For the encoding analysis, we estimated separate TRFs for each regressor (S1 and S2), and calculated the overall predictive power of the model at each MEG sensor. Guided by the results from the Selective Attention condition, for each participant we calculated the ‘TRF difference’ as the absolute difference between the two TRFs: |TRF(S1)-TRF(S2)|, within the 100-250ms time-window where significant effects of Selective Attention were observed and within the significant ROI. We then calculated the Pearson’s correlation coefficient r between the TRF-difference and the predictive power of the model, across all 100 different S1-S2 labelings, and tested whether the resulting correlation coefficients across all participants were significantly positive, using a two-tailed t-test. Similarly, for the decoding analysis we quantified the Pearson correlation between the reconstruction accuracies for S1 and S2 for each participant, across 100 different S1-S2 labelings, and compared the correlation coefficient to zero at the group level, using a two-tailed t-test.

Last, another advantage of the encoding searchlight analysis is that it allowed deriving a specific ‘best hypothesis’ as to the participants’ internal allocation of attention on a trial-by-trial basis. This ‘best hypothesis’ is defined as the S1-S2 labeling that yielded the highest predictive power, i.e., the one that provides the best estimation of the actual neural data for each participant (Fig 6b, shown in red). To test whether this ‘best hypothesis’ is a plausible proxy for the true allocation of attention under Distributed Attention, despite its ‘ground truth’ ambiguity, we compared it to the results obtained during Selective Attention. Specifically, we compared the predictive power as well as the absolute differences between the estimated TRFs for the two regressors in the Distributed Attention ‘best hypothesis’ vs. the Selective Attention condition, using a paired two–tailed t-test (averaged across the ROI sensors; and for the TRFs in a time window of 100-250ms). If the ‘best hypothesis’ explains the recorded neural responses to a similar degree as in the Selective Attention condition, this would indicate that the ‘best hypothesis’ is a viable hypothesis for the specific prioritization-scheme employed by each participant.

## Results

### Behavior

Accuracy on answering the comprehension questions was well above chance level in both conditions but were significantly better in the Selective Attention condition (88.58 ± 5.02%, mean±std) relative to the Distributed Attention condition (71.53 ± 8.67%, mean±std; two tailed paired t-test, t = 8.848, p < 10^−10^, Fig. 2a). In the Distributed Attention condition, we found no significant difference in accuracy on questions regarding the narrative asked about first (72.61 ± 9.95%) vs. second (70.45± 9.35%; mean±std; two tailed paired t-test, t = 0.822, p = 0.415, Fig. 2a, right inset), indicating no significant recency/primacy effects on task performance.

When testing whether performance on either task was better in individuals with high vs. low WMC (median split; Fig. 2b), we found no significant difference in the Selective Attention condition (z=-0.672, p=0.501, two tailed Wilcoxon ranksum test), however for Distributed Attention, performance was significantly better in the high WMC group (z=-2.311, p=0.021, two tailed Wilcoxon ranksum test).

### Selective Attention: Group level modulation of neural speech tracking

We examined the neural representation of the two speakers during the Selective Attention condition, using both an encoding and decoding multivariate model. Figure 3c (top panel) shows the topographical distribution of the predictive power averaged across participants, which indicates how well the encoding TRF model predicts the neural activity at each sensor. This distribution showed the common four-pole topography characteristic of auditory responses. Although predictive power values are relatively low (r∼0.03), this scale is in line with the values reported in previous similar studies (Ding and Simon 2012a; Zion Golumbic et al. 2013; Crosse et al. 2016), and was found to be significantly higher than chance in 30 sensors, indicated in Figure 3c, bottom panel (p<0.05 one sided, relative to a null-distribution of permutated data). These sensors were used as a Region of Interest (ROI) for subsequent analyses. Visual inspection of the TRF time courses for to-be-attended vs. task-irrelevant speech showed 3 prominent peaks and troughs at time lags ∼80, ∼120 and ∼200 ms. In line with previous findings, the TRFs were significantly modulated by Selective Attention, with larger peaks for the to-be-attended speaker compared to the task-irrelevant speaker between 120-140ms and between 160-240ms (p=0.002, cluster corrected over time, within the ROI sensors; Figure 3d).

In addition, we applied the decoding approach, which assesses the ability to reconstruct the acoustic envelopes of the to-be-attended and the task-irrelevant speech from the neural data. Here too, our group-level results replicate previous findings, demonstrating significantly higher reconstruction accuracy of the to-be-attended speaker (r = 0.07±0.02) vs. the to-be-ignored speaker (r = 0.05±0.03; mean±std; t = 4.044, p =0.00017 in a two-tailed paired t-test; Fig. 3e). Taken together, these results replicate previous findings regarding Selective Attention, indicating that the neural representation of two competing speakers is modulated by their task relevance, at least at the group level.

### Selective Attention: Individual level Selective-Attention Bias

We next turned to investigate the prevalence and size of this effect in individual participants. To this end, we calculated the Attention-Bias Index for each participant and normalized it to the distribution of bias values derived from 100 ‘attention-agnostic’ models (see Methods). This allowed us to estimate a normalized Attention-Bias Index in each participant and to ascertain whether it was significantly larger than could be achieved by-chance (Figure 4 a,b). Counter to our expectation, the normalized Attention-Bias indices varied substantially across participants, ranging from negative to positive values, with a mean normalized Bias Index of Z=1.18±1.53 (mean±std, 88^th^ %^tile^ of a standard normal distribution, Fig. 4c). Moreover, when looking at individual participants, the normalized Bias Index exceeded chance level only in 9 out of 27 (33%; Z > 1.96, examples shown on the left in Fig. 4b; all participants in Fig. 4c). For almost all other participants the normalized Bias Index fell within the range of the attention-agnostic null distribution, indicating that the difference in reconstruction accuracy of to-be-attended vs. task-irrelevant speech was not significantly larger than could be achieved by chance (see example on the right in Fig. 4a). One participant had a significantly negative normalized Attention-Bias Index (Z <-1.96).

In order to test whether the lack of an Attention-Bias in some participants merely reflects poor speech tracking overall and/or poor signal to noise, we compared the overall reconstruction accuracy in the two subgroups – participants with significant Attention-Bias (N=9) and those without it (N=18). However, we found no significant difference between them (one sided Wilcoxon ranksum test: z = 1.106, p=0.866, BF10 = 0.209 moderate evidence for the null; Figure 4c, left panel). This suggests that the participants who did not exhibit significant Attention-Bias, maintained equally-good neural representation for both speakers. We also found that these participants achieved similarly good performance on the Selective Attention task as the participants who did show significant Attention-Bias (one sided Wilcoxon ranksum test : z=1.424, p=0.923, BF10 = 0.188 moderate evidence for the null; Figure 4c right panel), indicating that neural Attention-Bias is not a prerequisite for adequately comprehending to-be-attended speech. Moreover, we did not find any correlation between WMC scores and the normalized neural Attention-Bias participants exhibited (Pearson correlation coefficient r=0.06, p=0.766) under Selective Attention.

### Distributed Attention: Neural speech tracking of equally task-relevant speakers

We wanted to test whether under Distributed Attention, where two speakers are both equally task-relevant, they are also equally represented in the brain response **(equal-representation hypothesis)**, or whether there is evidence from the neural data for a biased representation - either systematic or non-systematic - of one of the concurrent speakers **(biased-representation hypothesis)**. We first tested for systematic bias, either toward a particular speaker (Gender-Bias; preferring the female/male speaker) or towards a particular ear (Spatial-Bias; preferring speech presented from the right/left). At the group level, we did not find any significant Gender-or Spatial-Bias (Gender-Bias: t = 0.074, p=0.941; Spatial-Bias: t = 1.028, p=0.314, one sample t-test vs. 0; Fig. 5b and c, respectively). However, at the individual level, we found evidence for some form of systematic bias in 8/27 participants (∼30%). Of them, 4 participants showed a preference for one of the speakers (n=3 towards the female speaker; n=1 towards the male speaker; Fig. 5b, left lower panel, bold lines for Z>1.96), 3 participants showed a preference for one ear (n=2 towards the left ear; n=1 towards the right ear; Fig. 5c right lower panel, bold lines for Z>1.96), and 1 participant showed a preference for the male speaker as well as for the right ear.

Besides a systematic bias towards spatial location or speaker gender, we also tested for non-systematic biased-listening, in which one speaker/ear is preferentially represented in each trial. Since we do not have access to this internal ‘ground truth’, we used a searchlight approach in which 100 random S1-S2 labelings were tested. In the encoding version of this analysis, TRFs were estimated for S1 and S2 and the predictive power for the encoding model was estimated for each labeling (Fig. 6a,Fig. 3b). We then tested whether labelings that had higher predictive power (i.e., explained more variance in the neural data) generated similar TRFs for S1 and S2 (a pattern that would support the equal-representation hypothesis) or whether the TRF for one regressor was amplified, in line with the biased-representation hypothesis (Fig. 6b). We found the latter pattern, with a significantly positive correlation between the predictive power of different S1-S2 labelings and the differences between TRF amplitudes for S1 and S2 (correlation coefficient = 0.552 ± 0.110, mean ± std, p<10^−10^, t = 26.05, two-tailed t-test against 0; Fig. 7a). This suggests that the S1-S2 labeling with the best predictive power also yielded modulated TRFs for S1 vs. S2, in line with the biased-representation hypothesis.

**Figure 7.**
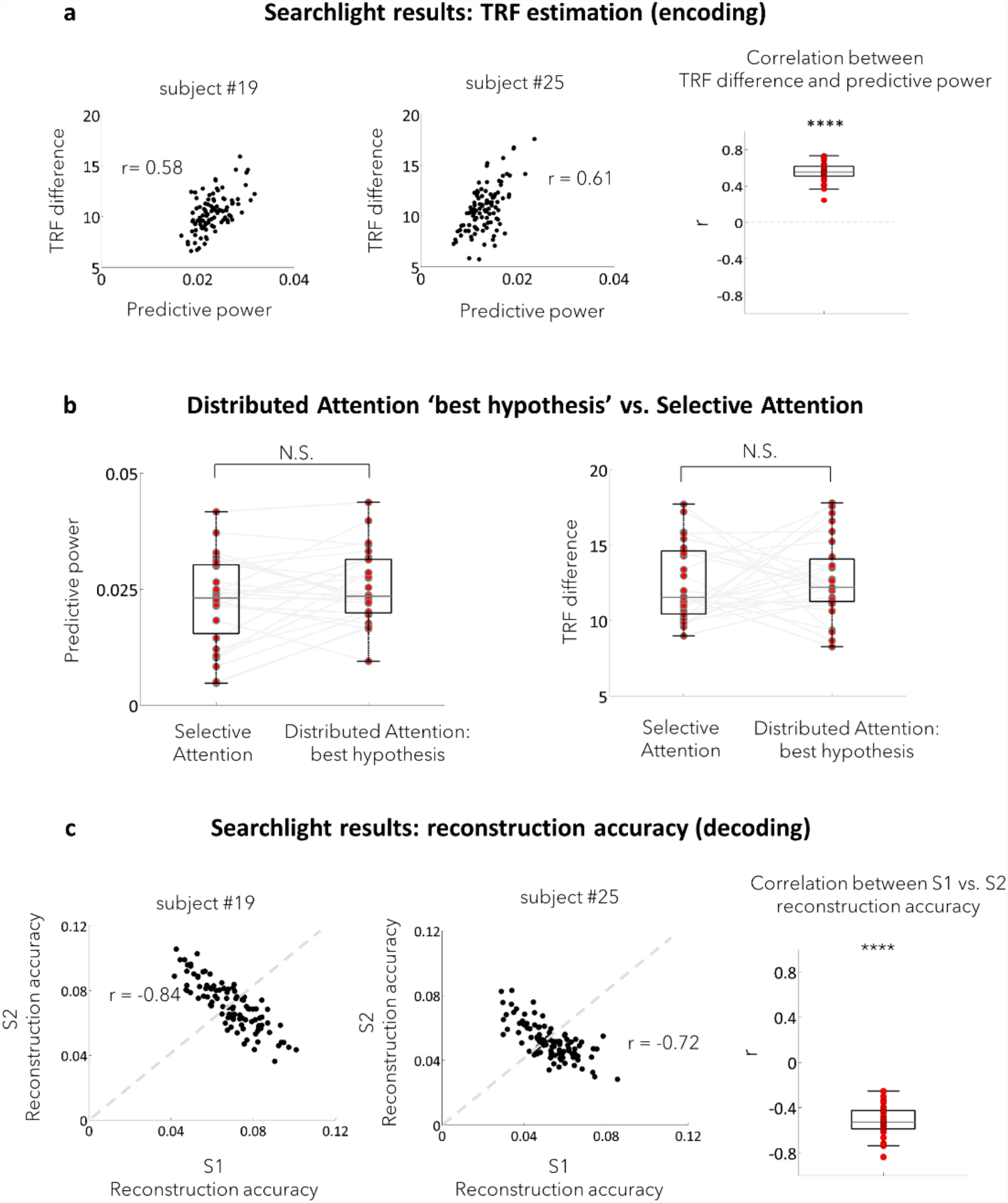
Distributed Attention: Searchlight results. **(a)** Encoding results. Left: Example from two participants, showing a positive correlation between the TRF difference for S1 and S2 (absolute difference) and the predictive power, across 100 different labelings (each dot represents a single S1-S2 labeling scheme). Both subjects demonstrate statistically significant positive correlations, indicating that the labelings that increasingly predict the neural data are those in which the estimated TRFs for S1 and S2 are increasingly different. Right: Distribution of the correlation coefficients (r) between the TRF difference and predictive power across S1-S2 labelings, for all participants. The correlation coefficient values were significantly positive at the group level (p<10^−10^), supporting the biased representation hypothesis. Red circles denote individual values and the box represents the 25-75^%tile^. **(b)** Theoretical predictions for the equal-representation (right) and biased-representation (left) hypotheses, for encoding results: Under the **biased-representation hypothesis**, we would expect a positive correlation between the difference in the TRFs estimated for S1 and S2 and the predictive power of the model, with the labeling schemes that best estimates the neural response (i.e., with the highest predictive power; ‘best hypothesis’ – indicated by a red dot) to also have the highest difference between the two estimated TRFs. Conversely under the **equal-representation hypothesis**, we would expect both narratives to be similarly represented in the neural response and therefore any differences between the TRFs are not expected to produce better predictions of the neural response. This would manifest in no significant correlation. **(c)** Theoretical predictions for the equal-representation (right) and biased-representation (left) hypotheses, for decoding results: Under the **biased-representation hypothesis**, we expect the reconstruction accuracies of the two streams to demonstrate a tradeoff, manifested in a negative correlation between the reconstruction of S1 and S2. Conversely, under the **equal-representation hypothesis**, we do not expect to find any correlation between the reconstruction accuracies of the two streams (or perhaps a positive correlation), as they are expected to be similarly represented in the neural response.

The S1-S2 labeling scheme that yielded the highest predictive power in each participant was considered to be the ‘best hypothesis’ for how they may have prioritized among the two speakers on a trial-by-trial basis in the Distributed Attention condition. We directly compared the predictive power and the differences in the estimated TRFs for S1 and S2 under this ‘best hypothesis’ labeling vs. those obtained in the Selective Attention condition. In both comparisons we found no significant differences (predictive power: t=-1.770, p=0.088; TRF differences: t=-0.428, p=0.672, two-tailed paired t-test, Figure 7b), which suggests that the Distributed Attention ‘best hypothesis’ yields a similar modulation of TRFs as is found during Selective Attention (i.e., enhanced TRF for one regressor), and explains the recorded neural signal to a similar degree.

Finally, to complete these results, in the decoding searchlight approach, we calculated the reconstruction accuracies for S1 and S2 [r(S1) and r(S2)] across 100 different labeling schemes. We then tested the correlation between r(S1) and r(S2) across different labelings, and found a highly significant negative correlation (−0.523± 0.139 mean±std, t = -19.544, p<10^^-10^, two-tailed t-test against 0; Figure 7c). This pattern suggests there is a tradeoff in the degree to which the two competing speakers are represented in the neural signal (Figure 6c, left). Taken together, the converging results of both the decoding and encoding analyses are more compatible with the biased-representation hypothesis, suggesting that during our Distributed Attention task listeners internally prioritized one of the speakers in each trial, and did not (or could not) represent both speaker to the same degree.

## Discussion

This study offers unique insight into the intersection between a listener’s behavioral goals and the system’s inherent capacities and limitations for processing competing speech. The emerging pattern from comparing Selective and Distributed Attention suggests that the observed neural speech tracking response to competing speech does not always directly reflect their instructed task-relevance. On the one hand, we show that despite the well-documented enhanced representation of to-be-attended speech during Selective Attention (Hillyard et al. 1973; Hansen and Hillyard 1983; Woods et al. 1984; Bidet-Caulet et al. 2007; Manting et al. 2020), this selection is not exclusive and task-irrelevant speech is also encoded quite well (sometimes just as well as to-be-attended speech). On the other hand, we also show that during Distributed Attention concurrent speech are not represented equally, even though this would be most beneficial for the task. Rather, we find evidence for a tradeoff in the neural representation of these equi-task-relevant speech, which likely reflects inherent bottlenecks for fully processing more than one speech stream (Broadbent 1958; Treisman 1964; Duncan 1980; Lachter 2004; Bronkhorst 2015; Kawashima and Sato 2015; White et al. 2019).

In interpreting the results of our study, it is important to bear in mind that the neural metric of speech tracking used here (envelope-following response) primarily captures the acoustic representations of input in auditory cortex (Ding and Simon 2012a; Mesgarani and Chang 2012; Zion Golumbic et al. 2013; Fiedler et al. 2019a; Har-shai Yahav and Zion Golumbic 2021). Therefore, the current results should not be taken as an exhaustive description of all the ways in which attention may affect neural processing of speech. Rather, the neural speech tracking metric affords a more specific lens into the downstream modulatory effects that top-down attention and/or the deployment of different listening strategies has on the bottom-up sensory encoding of natural speech.

### Modulation of speech tracking by Selective Attention is not ubiquitous

In the Selective Attention condition, we replicated the well-established bias towards to-be-attended speech, which manifests as an enhanced speech tracking response (TRF) and better reconstruction accuracy for to-be-attended vs. task-irrelevant speech (Kerlin et al. 2010; Ding and Simon 2012a, 2012b; Mesgarani and Chang 2012; Power et al. 2012; Zion Golumbic et al. 2013; O’Sullivan et al. 2015; Fuglsang et al. 2017; Fiedler et al. 2019). However, despite being highly significant at the group-level, this effect was not ubiquitous across all participants, and was statistically significant only in a subset of participants (∼30%). To the best of our knowledge, previous studies have not directly addressed this inter-subject variance in attentional-bias, however several pervious papers that present individual-level data seem to exhibit similar variability (Ding and Simon 2012a, 2012b; Fuglsang et al. 2017; Rosenkranz et al. 2021). In the current data set, inter-subject variability in the magnitude of attentional bias was not correlated with working memory capacity (OSPAN), which sometimes drives individual differences in attentional abilities (Beaman et al. 2007; Sörqvist and Rönnberg 2014a; Lambez et al. 2020). Importantly, in our data, the overall speech-reconstruction accuracy was comparable in both sub-groups, indicating that the lack of attentional bias in some participants was not simply due to reduced signal to noise. Moreover, both sub-groups performed equally well on the behavioural task. Therefore, we do not interpret these data as suggesting that participants who did not exhibit significant bias in their neural response towards to-be-attended speech simply did not attend selectively to the prescribed speaker. Rather, we believe they suggest that there is no necessary-dependency between ‘internal allocation of attention’ and the modulation of sensory responses to competing speech, even though these often occur together. Below we offer two possible explanations for what this inter-subject variability might reflect, other than inappropriate allocation of attention.

One possibility, inspired by ‘Load Theory of Attention’ (Forster and Lavie 2008; Murphy et al. 2017) is that the Selective Attention task used here was sufficiently easy (answer comprehension questions), and the perceptual competition between stimuli was relatively moderate (with high spectral and spatial separation between speakers), such that residual processing resources were available to encode task-irrelevant stimuli. According to this perspective, achieving good performance on a Selective Attention task does not necessarily require attention-biasing of sensory level neural responses, particularly under conditions of moderate perceptual and cognitive loads. Alternatively, the inter-subject variability may be related to the speech tracking measures themselves and what they represent. Specifically, it is possible that even though in some participants the envelope of to-be-attended and task-irrelevant speech could be reconstructed just as well from the neural signal, the underlying neural code for each speaker was different. This possibility is supported by previous studies showing that decoders trained separately on to-be-attended and task-irrelevant speech achieved good reconstruction accuracies for both, but not when the to-be-attended decoder was used to predict task-irrelevant speech and vice versa (Ding and Simon 2012a; O’Sullivan et al. 2015). These two interpretations are not mutually exclusive and, importantly, offer specific hypotheses regarding the neural implementation of Selective Attention, that go beyond the simple amplification of to-be-attended speech. Since studying individual differences was not the explicit goal of this study, but rather a post-hoc observation from the data, we encourage future research to take a closer look at these hypotheses, as well as at how effects of attention on the speech tracking response manifest at the individual level.

### Biased listening strategy for achieving Distributed Attention

In comparison with Selective Attention, where there is a single focus of attention, Distributed Attention is a substantially more difficult task, that requires extensive top-down control for allocating limited linguistic processing resources among several stimuli (Treisman 1964; Duncan 1980; Koelewijn et al. 2012; Bronkhorst 2015; Kawashima and Sato 2015; Mccloy and Lee 2015; Gagné et al. 2017; Lambez et al. 2020; Yuriko Santos Kawata et al. 2020; Agmon et al. 2021). The more taxing nature of Distributed Attention was confirmed by our behavioral results, which showed significantly reduced accuracy in answering questions about two concurrent narratives relative to one (in the Selective Attention condition). Moreover, performance on the Distributed Attention task was better in individuals with high WMC, supporting its reliance on the availability of more executive processing resources and / or the more effective management of these resources (Beaman 2004; Koelewijn et al. 2012; Tsuchida et al. 2012; Sörqvist and Rönnberg 2014; Lavie et al. 2014; Rudner 2016; Wiemers and Redick 2018; Lambez et al. 2020).

Despite its more demanding nature, participants still performed the Distributed Attention task relatively well, which prompted us to investigate how the two speech stimuli were encoded to achieve this level of performance. Conducting speech tracking analysis of the neural activity in the Distributed Attention condition posed a unique methodological challenge, since there was no a-priori “correct” way to label the two concurrent speech-stimuli as two separate regressors (S1 and S2), as is typically done for Selective Attention. Nonetheless, by testing a range of different possible S1-S2 labeling schemes we were able to investigate whether there was evidence for systematic and/or non-systematic bias in the neural representation of two equi-task relevant speakers. In this approach, each labeling scheme represents a hypothetical listening strategy in which one of the two competing speakers was preferred in each trial. If we had found similar predictive-power (in the encoding model) and similar reconstruction-accuracies (in the decoding model) across different labeling schemes, this would have indicated that allocation of speakers to S1/S2 was fully interchangeable, supporting the ‘equal-representation’ hypothesis. However, this was not what we found.

Rather, our results indicate that some individuals (8/27, ∼30%) showed systematic biased listening, with a preferences for a particular speaker or spatial location. In addition, both our encoding and decoding analysis showed evidence for a tradeoff in the neural representation of the two concurrent speakers, on a non-systematic, trial-by-trial basis. Specifically, the encoding analysis showed that the labelling schemes which explained the most variability in the neural data (“best hypothesis”) were those in which there were larger differences in the speech tracking TRFs for the two speakers. Interestingly, the predictive power of this “best hypothesis” labelling scheme was just as good as found in the Selective Attention case, further validating its candidacy as a possible listening strategy the was actually used. In addition, the magnitude of the TRF differences under this “best hypothesis” was similar to that found between to-be-attended and task-irrelevant speech under Selective Attention. Therefore, despite the inherent ambiguity regarding participants’ internal attention state/preferences, this pattern suggests that even under Distributed Attention, individuals employ some form of biased listening among the two competing speech.

This was also supported by the decoding results, where reconstruction accuracies for the two speakers across different labeling schemes were negatively correlated, such that in schemes where high reconstruction accuracy was obtained for one regressor (e.g., S1), the reconstruction accuracy for the other regressor (e.g. S2) was low. Taken together, this pattern of results does not support the ‘equal-representation’ hypothesis, but rather is more in line with the ‘biased-representation’ hypothesis, which posits that even when attempting to divide attention among two speakers, one speaker is nonetheless preferentially represented over the other, plausibly due to inherent bottlenecks in linguistic processing (Broadbent 1958; Treisman 1964; Duncan 1980; Lachter 2004; Bronkhorst 2015; Kawashima and Sato 2015; White et al. 2019).

It is important to stress that our findings supporting ‘biased-representation’ do not imply that *only* one speaker was selected for processing in each trial (as might be the case in Selective Attention). Rather, it indicates that one speaker was tracked and encoded *more accurately than the other*, throughout the course of each trial. This preference could manifest either as dynamic shifts of attention between the two speakers, with overall more time spent attending to one vs. the other (‘glimpsing’; Vestergaard et al. 2011; Fogerty et al. 2018; Shavit-Cohen and Zion Golumbic 2019; Spadone et al. 2021), or as a constant preference throughout the trial to one speaker (akin to Selective Attention). Indeed, several limiting factors preclude providing a more detailed account of how this biased listening strategy is employed from the current study. One limitation is our inability to reliably characterize the within-trial dynamics of Distributed Attention, and hence test whether biased listening is achieved through shift of attention between speakers (‘glimpsing’). This is due to the insufficient temporal resolution of the speech tracking measure, coupled with the ‘ground truth’ problem, i.e. the lack of an ability to validate the listeners’ locus of attention on a moment-to-moment basis. Another related limitation is the low sensitivity of the behavioral task - answering 3 comprehension questions about each narrative - which only provides a gross behavioral readout of comprehension and lack sufficient temporal resolution for capturing attention-shifts and/or tradeoff between speakers. This is an inherent limitation of many similar studies of speech-processing, which strive to limit the interruption of natural listening with a secondary task (e.g. detect targets), and also wish to avoid extensive post-stimulus questioning. Development of an alternative behavioral test in future studies would be extremely valuable, particularly for testing the behavioral-relevance of the Distributed Attention ‘best hypothesis’ identified through our searchlight analysis, paving the way towards overcoming the ‘ground truth’ problem of attention-studies. That said, the task used here does capture the level of speech comprehension that we generally aim to achieve in real-life contexts. As such it provides a useful indication of how well individuals deal with the presence of task-irrelevant speech (Selective Attention) or the need to monitor two speakers at once (Distributed Attention), in order to achieve an ecologically-relevant level of understanding.

## Conclusions

By comparing two extreme cases of prescribed attention – Selective vs. Distributed – we show that the neural encoding of concurrent speech is not always modulated as might be expected by the task-instructions only. These findings encourage us to adopt a more nuanced perspective when thinking about the challenge of paying attention in multi-speaker contexts. Rather than assuming discrete categorization of speech-stimuli as “attended” or “ignored”, individuals likely employ a dynamic continuum of prioritization, where the degree to which concurrent speech is processed is affected not only by the listeners explicit behavioral goals but also by the limits/excess of available resources (Forster and Lavie 2008; Murphy et al. 2017) as well as contextual factors (Kaya and Elhilali 2017). As such, Selective and Distributed Attention to speech can be viewed as two extremes of this continuum, which - perhaps not surprisingly - recent studies have shown engage a largely overlapping network of temporal-frontal-parietal brain regions (Lipschutz et al. 2002; Hill and Miller 2010; Miller 2015; Moisala et al. 2015; Yuriko Santos Kawata et al. 2020; Agmon et al. 2021). As research of attention to speech progresses towards studying performance in increasingly ecological contexts (Fiedler et al. 2019; Shavit-Cohen and Zion Golumbic 2019, Hölle et al., 2021), understanding the systems’ flexibility to accommodate concurrent speech, and the underlying moment-to-moment dynamics, will become even more important.

